# Quantum aspects of evolution: a contribution toward evolutionary explorations of genotype networks via quantum walks

**DOI:** 10.1101/2020.07.10.197657

**Authors:** Diego Santiago-Alarcon, Horacio Tapia-McClung, Sergio Lerma-Hernández, Salvador E. Venegas-Andraca

## Abstract

Quantum biology seeks to explain biological phenomena via quantum mechanisms, such as enzyme reaction rates via tunneling and photosynthesis energy efficiency via coherent superposition of states. However, less effort has been devoted to study the role of quantum mechanisms in biological evolution. In this paper, we used transcription factor networks with two and four different phenotypes, and used classical random walks (CRW) and quantum walks (QW) to compare network search behavior and efficiency at finding novel phenotypes between CRW and QW. In the network with two phenotypes, at temporal scales comparable to decoherence time T_D_, QW are as efficient as CRW at finding new phenotypes. In the case of the network with four phenotypes, the QW had a higher probability of mutating to a novel phenotype than the CRW, regardless of the number of mutational steps (i.e., 1, 2 or 3) away from the new phenotype. Before quantum decoherence, the QW probabilities become higher turning the QW effectively more efficient than CRW at finding novel phenotypes under different starting conditions. Thus, our results warrant further exploration of the QW under more realistic network scenarios (i.e., larger genotype networks) in both closed and open systems (e.g., by considering Lindblad terms).

## Background

Quantum biology is a novel discipline that uses quantum mechanics to better describe and understand biological phenomena (Mohseni 2014; Brookes 2017; McFadden and Al-Khalili 2018). Over the last 15 years, there have been theoretical developments and experimental verifications of quantum biological phenomena (McFadden and Al-Kahlili 2014; Brookes 2017) such as quantum tunneling effects for the efficient workings of enzymes at accelerating biological metabolic processes (e.g., Klinman and Cohen 2013), and quantum superposition for efficient energy transfer in photosynthesis (Panitchayangkoon et al. 2010). The area of quantum evolution (McFadden and Al-Kahlili 1999), in which it is suggested that DNA base pairs remain in a superposition by sharing the proton of hydrogen bonds, still remains speculative and has practically stagnated since its theoretical inception twenty years ago (Ogryzko 1997; McFadden and Al-Kahlili 1999). However, recent theoretical developments on quantum genes (e.g., Brovarets’ and Hovorun 2015) suggest that further exploration of the superposition mechanism in evolution is worth undertaking.

### Theoretical framework: a) quantum measurement device

From biological principles, genes do not vary in a continuous fashion, they are digital objects (i.e., a sequence of discrete nucleotides); such discontinuity renders mutations as quantum jumps between different states or possible variations of a gene (Schrödinger 1944; Godbeer et al. 2015). In other words, genes function as discrete packets, which are akin to quantum digital objects over which computations are performed (Lloyd 2008). Hence, the theoretical framework of quantum mechanics offers two characteristics that are fundamental for life and its evolution: digitalization and probabilistic variation among the discrete states a quantum system can take (e.g., DNA nucleotides; Lloyd 2008).

Focusing on DNA, the genetic code is ultimately determined by hydrogen bonds of protons shared between purine and pyrimidine nucleotide bases (McFadden and Al-Khalili 1999). Nucleotides have alternative forms knows as tautomers, where the positions of the hydrogen protons in the nucleotide are swapped, changing nucleotides chemical properties and affinities (Watson and Crick 1953a,b). Such changes make the DNA polymerase enzyme to sometimes pair wrong nucleotides (e.g., a tautomeric thymine with a guanine), generating mutations that change the genetic information and possibly the encoded protein (McFadden and Al-Khalili 1999, 2014; Fig. 1). An important consequence of this process, since genes can be thought of as quantum systems, is that nucleotides’ hydrogen bridges can be described as a quantum superposition, where protons can be found at both sides of the DNA chain at the same time (i.e., the physical variable in a superposition is the hydrogen proton joining DNA nucleotides; quantum genes), hence allowing the system to be described by a wave function (McFadden and Al-Kahlili 1999, Godbeer et al. 2015). A measurement (e.g., a chemical, UV light from the environment) can collapse the wave function producing either a normal base pair or a mutation (McFadden and Al-Kahlili 1999, 2014). Thus, quantum processes can be of relevance in the generation of mutations (i.e., adaptive mutations) when influenced by the surrounding environment (i.e., selective factors; Brovarets’ and Hovorun 2015; Godbeer et al. 2015; Fig. 2), playing an important role in the exploration of evolutionary space (e.g., n-dimensional genotype networks, as introduced shortly in this manuscript).

**Fig 1.**
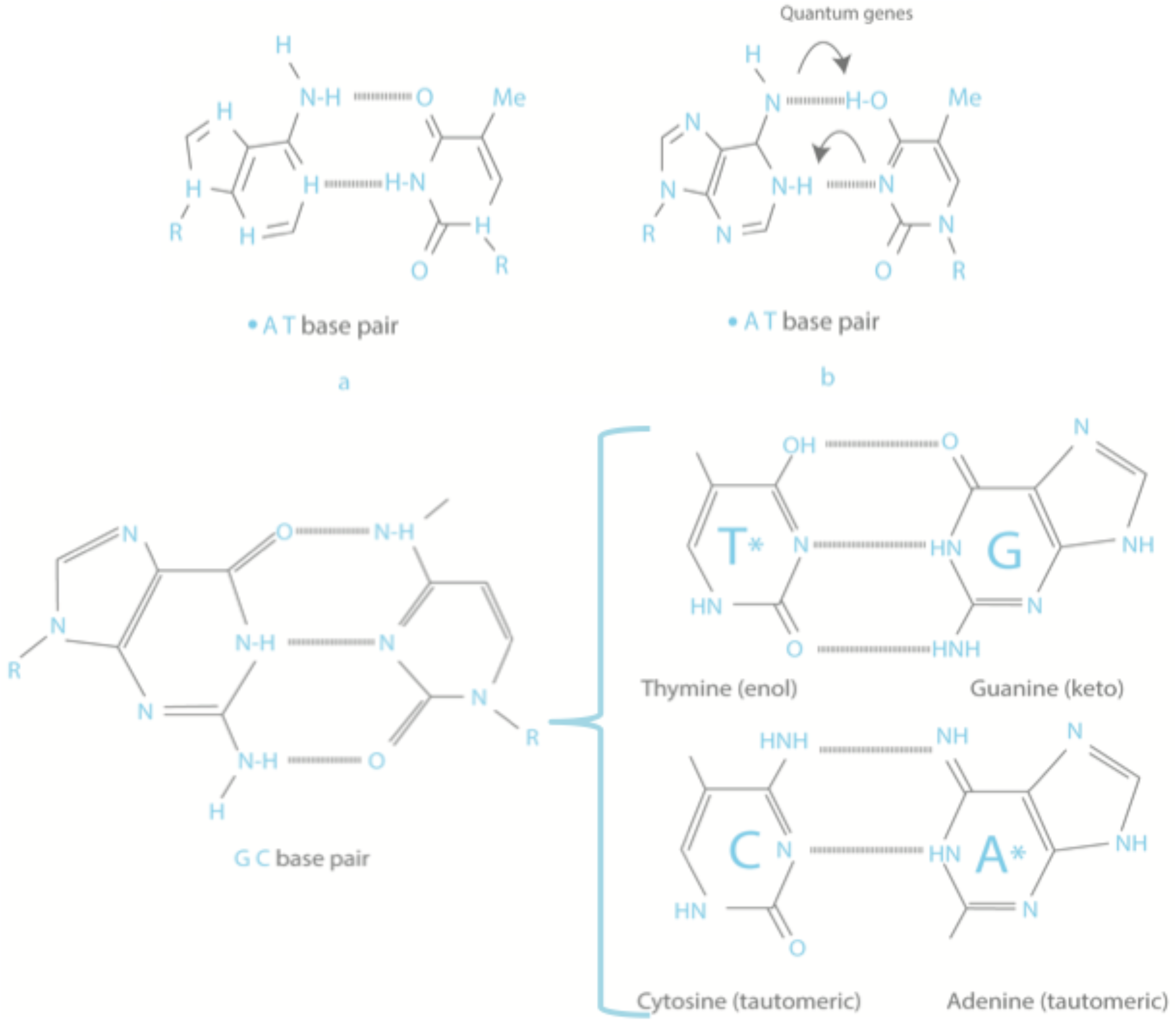
At the top, (a) shows a correct A-T base pairing, whereas (b) shows an A-T base pair with their hydrogen protons switched. At the bottom, on the left a correct G-C base pair and on the right two tautomeric base pairs (modified from McFadden and Al-Kahlili 2014).

**Fig 2.**
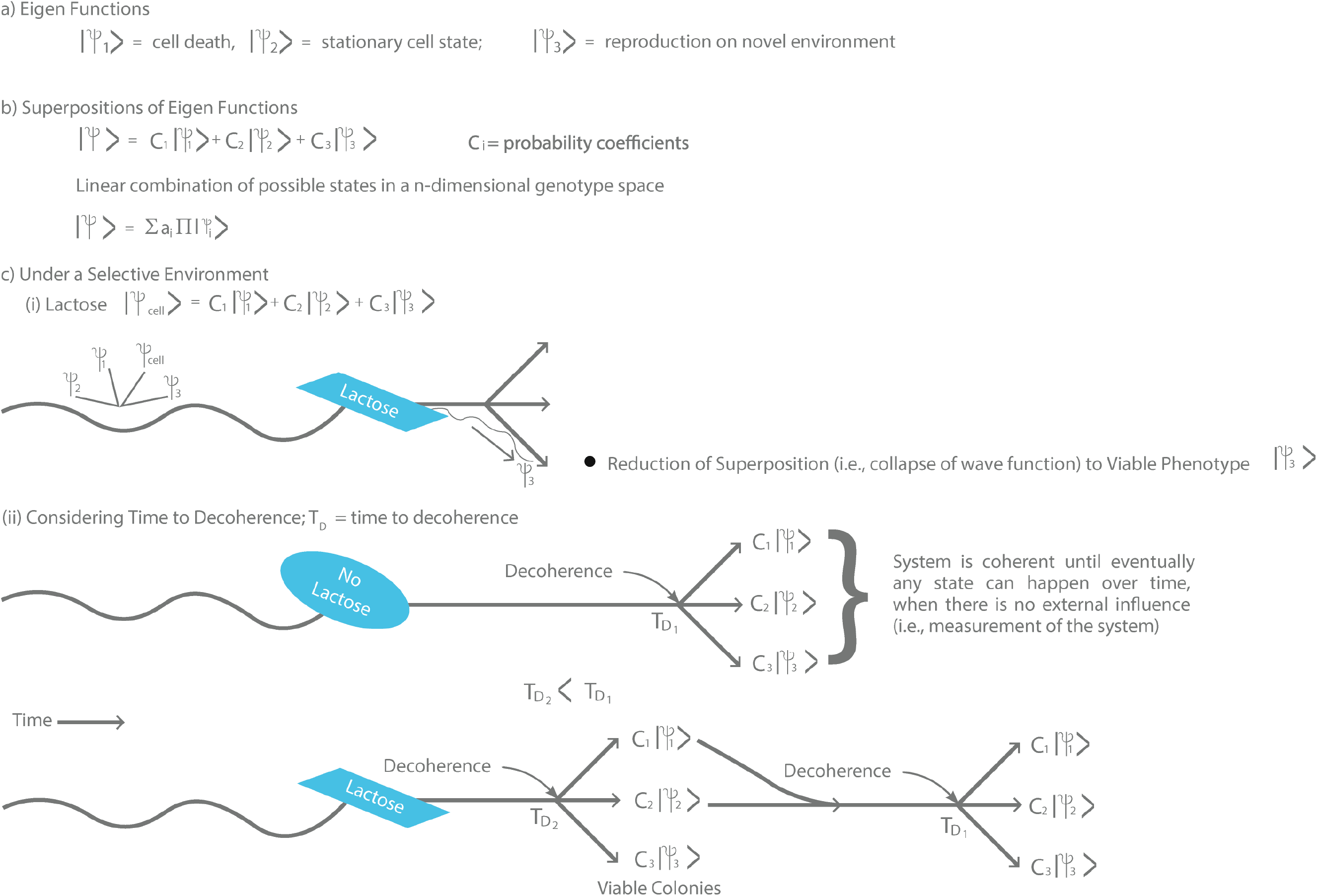
Decoherence process of a quantum wave function with three possible states under the influence of an environmental factor (measurement). **a)** Three possible bacterial cell states (we only use three for simplicity, but it can include all n-dimensional neighbors in a genotype network, see Fig. 4), represented by state vectors. **b)** Superposition of the three state vectors, which results in a linear combination of eigen functions each with a probability C_i_. **c)** The quantum wave function collapses toward the fit cell variant (i). When measuring time to decoherence (ii) all states of the quantum superposition under no selective conditions (i.e., no lactose) are indistinguishable by the bacterial cell, eventually collapsing to any of the possible states at T_D1_. However, when the environmental factor is present (i.e., lactose present) the time to decoherence will be shorter (T_D2_) and biased toward fit variants (i.e., adaptive mutation) able to grow and reproduce. Those bacterial cells that do not reduce toward the adaptive state, will remain in a quantum superposition. Thus, the quantum superposition will collapse to the adaptive state with higher probability under the environmental adaptive conditions (i.e., lactose present) compared to the time it takes to appear under non-selective environments (T_D2_ < T_D1_).

### Theoretical framework: b) n-dimensional genotype networks

A theory based on n-dimensional genotype space at different levels of biological organization (e.g., metabolism, gene regulation) has been developed to understand the evolution of innovations (Wagner 2011, 2014). A genotype network implies the existence of a vast connected network of genotypes (nodes in a network) that produces the same phenotype (Schuster et al. 1994). Genotypes in a genotype network can share little similarity (e.g., lower than 25%) and still produce the same phenotype (Wagner 2011). To understand the concept of a genotype network we will focus on metabolic reactions (Wagner 2011; Fig. 3).

**Fig 3.**
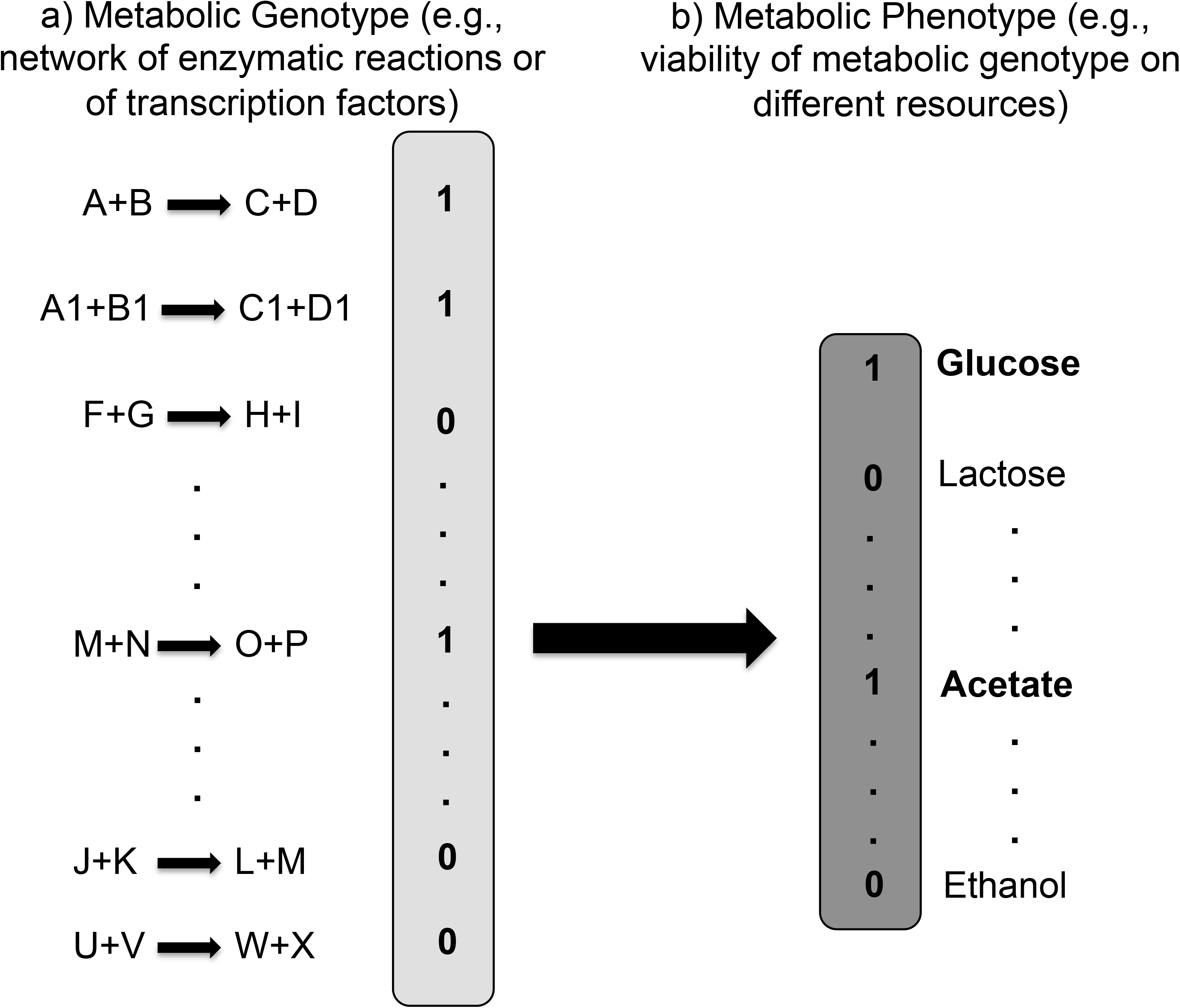
a) A list of metabolic reactions, a 1 next to a reaction indicates that an organism has such a metabolic path otherwise there is a 0. b) A list of resources that can be used (1) or not (0) by a metabolic genotype in order to synthesize all required biomolecules (see the text for details; modified from Wagner 2011).

A metabolic genotype is the total amount of chemical reactions that can be performed by the enzymes synthesized by an organism’s genotype (Wagner 2011). If we use digital (i.e., binary) categorization, then we can classify a metabolic genotype as a string of binary flags, indicating if the genotype has the information to synthesize a product that performs a metabolic reaction (represented by 1) or not (represented by 0; see Fig. 3). From current information we know there are about 10^4^ metabolic reactions (no organism can perform all of them; Samal et al. 2010), in which case we would have in binary space with 2^10,000^ different possible metabolic genotypes, which is a large universe of possibilities available for evolution to explore (Samal et al. 2010, 2011; Wagner 2014). Hence, the genetic space of metabolic genotypes is composed of all possible binary strings of length 10^4^, in this case a total of 2^10,000^. A way to measure differences between two metabolic genotypes in this vast space is to use the fraction of reactions that are not catalyzed by one genotype in reference to the other; the letter *D* represents such a measure (Rodrigues and Wagner 2009). The maximal value *D* = 1 would be achieved when the two metabolic genotypes do not have any reaction in common and *D* = 0 when they have identical metabolic genotypes (i.e., they would encode the same products or enzymes). Two metabolic genotypes would be neighbors if they differ only by a single reaction (a 1 in our binary coding of metabolic genotypes). Hence, the neighborhood of a metabolic genotype is composed by all those genotypes that differ by exactly one reaction from it; there would be as many neighbors as there are metabolic reactions (Fig. 4). Considering the different possible metabolisms one step away from a focal one, each neighborhood would be a large collection of metabolic genotypes organized in a hyper-dimensional cube. With this we build an n-dimensional network, where each genotype is a node in the network and the edges represent mutational steps, nodes connected by an edge differ exactly by one mutation (Rodrigues and Wagner 2009; Wagner 2014; Fig. 4).

**Fig 4.**
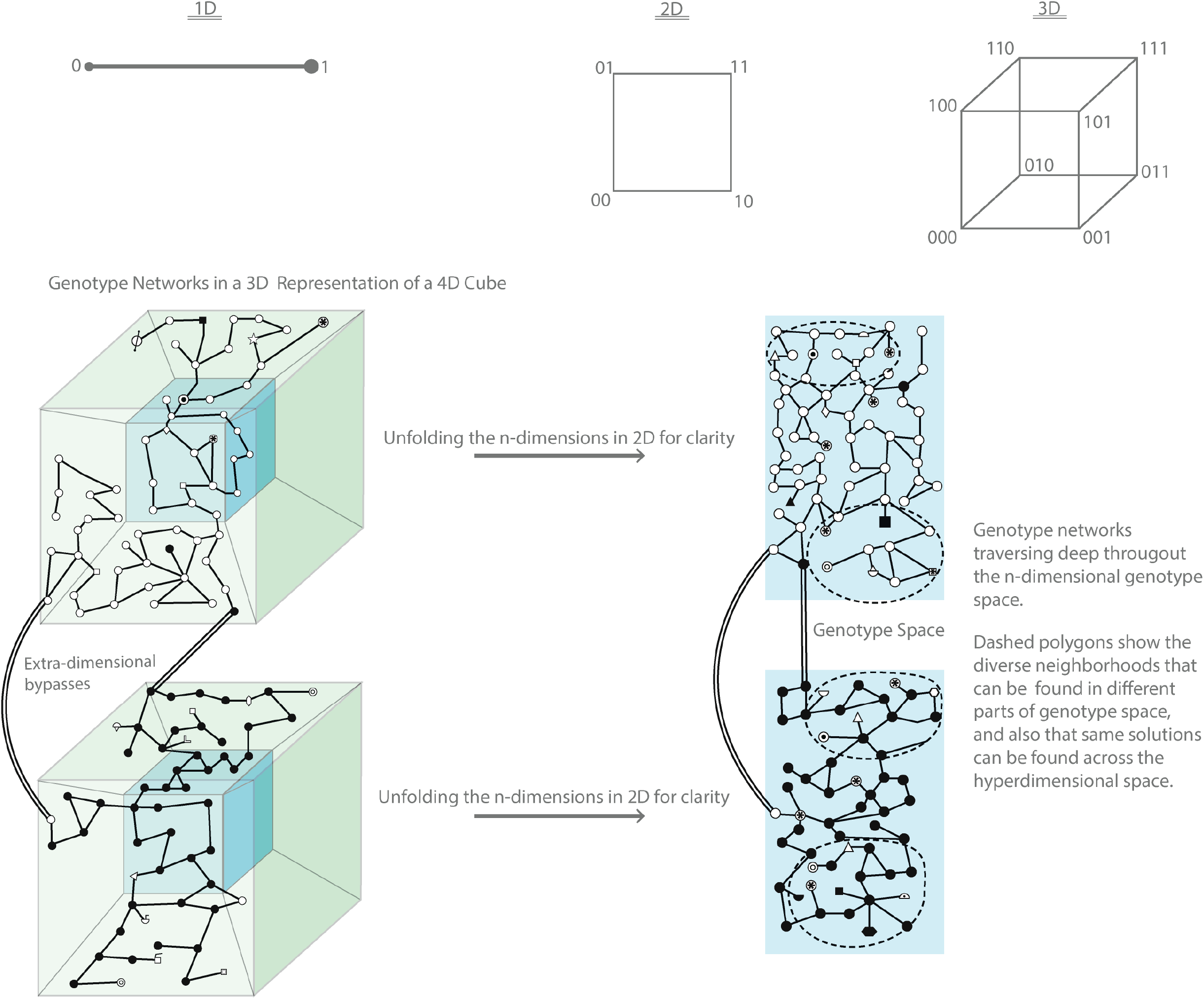
Representation of metabolic genotypes and phenotypes in different dimensions (modified from Wagner 2011). Networks in one, two, and three dimensions, where vertices are labeled with the binary strings that correspond to each dimension (1 = presence of metabolic pathway, 0 = absence of metabolic pathway; see Fig. 3). Two versions of a 3D representation of a four-dimensional cube are shown (i.e., the shadow of a Tesseract), each with its own section of a genotype network (i.e., network of white circles in upper panel and network of black circles in lower panel representing different phenotypes). Each line (i.e., link or edge) connecting two symbols represents a single mutational step. Genotype networks (those with same symbols) are vast across hyper-dimensions (genotype space), maintaining the same phenotype (i.e., robust to mutational changes across the network) even if genotype similarity is low (e.g., nodes on opposite sides of the genotype network). On the right side, we unfold the 4D cubes into 2D images for clarity. There, neighborhoods at different places of the genotype space are very diverse (different symbols inside dashed circles), which opens opportunities to find novel phenotypes. Some of the same evolutionary novelties can also be found at different neighborhoods, allowing for convergence. Each genotype network is connected to an n-number of other genotype networks via extra-dimensional bypasses (black double lines connecting genotype networks belonging to different phenotypic networks).

A metabolic phenotype is represented by all the environmental energy sources (e.g., glucose, methane) that can be used by a metabolic genotype to synthesize all biomolecules (e.g., amino acids, nucleotides) required for survival (Fig. 3). The metabolic phenotype can also be categorized as a binary string, a 1 represents a genotype network that can synthesize all required biomolecules relying solely on that specific source and a 0 otherwise; a phenotype with multiple ones means a metabolism that can produce all needed elements from many different sources (Wagner 2011). To calculate the number of possible phenotypes, we do the same as for metabolic genotypes; we raise two to the power of all the known different energy sources available. The set of those metabolic genotypes that have the same phenotype is what constitutes a genotype network. It has been shown computationally that similar (i.e., neighbors), as well as very dissimilar genotypes (as different as 80% of their metabolic reactions), can still preserve the same phenotype, demonstrating that genotype networks are plastic and robust (e.g., Wagner 2008; Rodrigues and Wagner 2011). This is a good feature for evolving populations because browsing the vast genotypic space becomes feasible and moderately free of risk (Rodrigues and Wagner 2009, Samal et al. 2010). However, how can new features evolve when a vast exploration leads us to the same viable result or phenotype? When comparing the neighborhoods of thousands of pairs of metabolic genotypes that are able to use the same energy source (i.e., they belong to the same phenotype network), but that are otherwise very different, it turns out that their neighborhoods are very different and diverse (i.e., novel phenotypes in one neighborhood might not be present in other neighborhoods of the same genotype network, Wagner 2014; Fig. 4). As the number of changed metabolic reactions increases, so does the number of unique phenotypes in a neighborhood, opening a bounty of novel phenotypes to an evolving population (Rodrigues and Wagner 2009). Furthermore, when comparing two genotype networks (i.e., networks that produce different phenotypes), the distance in genotype space that needs to be traversed to find a novel phenotype is rather small (i.e., the number of edges or mutational steps in the network separating nodes or genotypes with different phenotypes), raising the odds of finding novel traits (Wagner 2014, Rodrigues and Wagner 2011; Fig. 4). More impressive yet is the fact that networks other than metabolism, such as transcriptional regulatory circuits (Ciliberti et al. 2007, Espinosa-Soto et al. 2011) and the development of novel molecules (Li et al. 1996, Cui et al. 2002, Bastolla et al. 2003, Sumedha et al. 2007) have the same basic structure (Wagner 2011).

There is no true randomness as originally conceived in Darwinian evolutionary theory (e.g., Cairns et al. 1988; Hall 1995, 1997; Wagner 2012a). A series of experiments have shown that mutations are not completely random and that they can actually happen as a response to an environmental factor (e.g., Cairns et al. 1988, Rosenberg et al. 1994; Hall 1997, 1998; Hendrickson et al. 2002; Stumpf et al. 2007; Braun and David 2011; Livnat 2013). Thus, we are ultimately interested in the potential effect that specific environmental conditions (i.e., probing agents that collapse the quantum superposition) have on the proposed genetic quantum system and the evolutionary pathway followed under such conditions (e.g., Fig. 5). Yet, we must first understand how quantum processes behave under non-selective (i.e., neutral and in closed systems) scenarios, so we can determine their relevance for evolution. Thus, in this paper we explore how fast a quantum walk (QW) could explore an n-dimensional genotype network, *sensu* Wagner 2011 (i.e., a state space) and compare its performance with that of a classical random walk (CRW) (e.g., Farhi and Gutmann 1998). Then, we explore under what scenarios of the state space (i.e., mutational steps between different phenotypes) may the quantum process be more efficient than the classical one at finding novel states (i.e., phenotypes) in n-dimensional genotype networks (Wagner 2014; Aguilar-Rodríguez et al., 2017). That is, we provide proof of concept that genotype networks are the evolutionary fabric on which the earlier proposed quantum wave function (Ogryzko 1997; McFadden and Al-Khalili 1999) can operate, and then how the quantum wave function actually operates on such evolutionary fabric.

**Fig 5.**
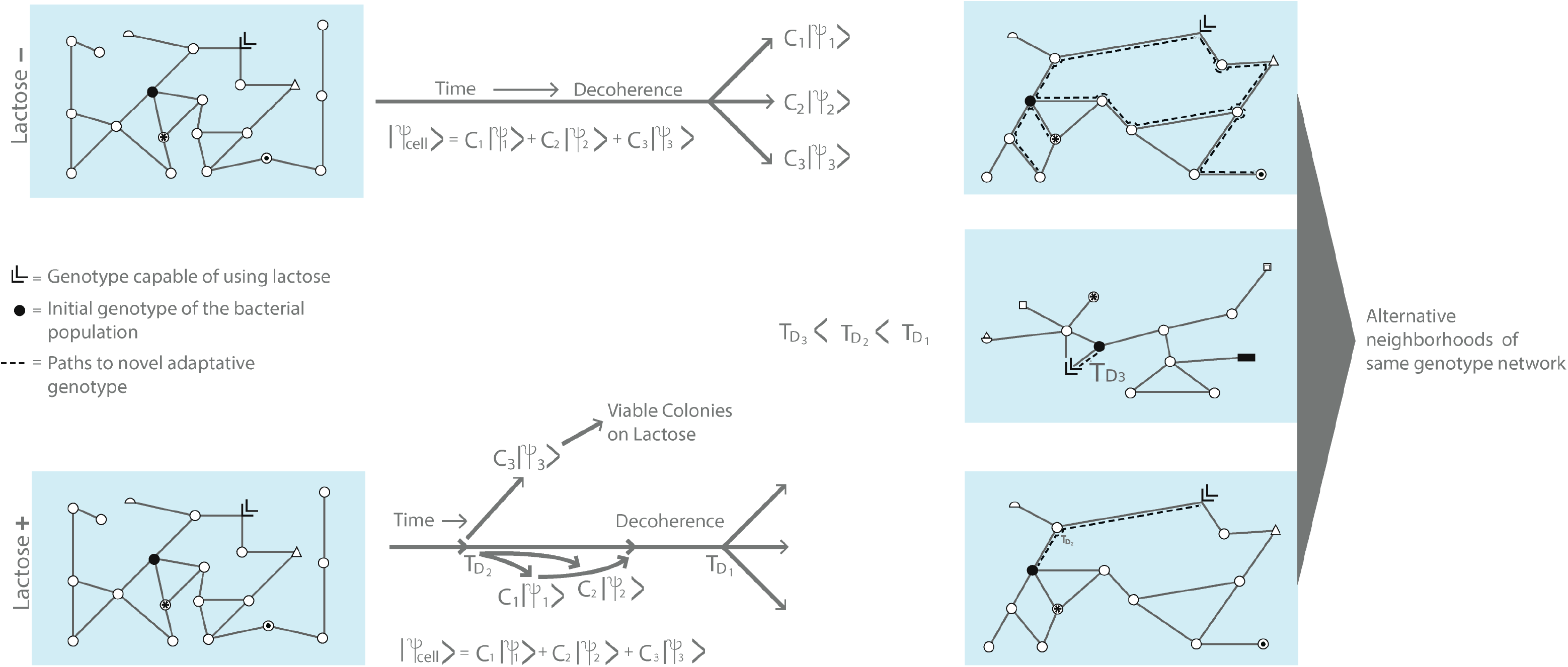
A bacterial genotype network under two environments without lactose (top) and with lactose (bottom). The superpositions of three possible cell states and times to decoherence are depicted in the middle, to the right of each genotype network (see Fig 2 for details). On the right hand side, there are three alternative neighborhoods of the original genotype network shown on the left. Different decoherence times (T_D_) to reach the genotype capable of using lactose are illustrated, based on different paths followed on different neighborhoods of the genotype network. The time to decoherence from the middle network on the right hand side is shorter compared to the other two (i.e., T_D3_ < T_D2_ < T_D1_).

## Methods

QW are more efficient at exploring one dimension (e.g., linear) and two dimension (e.g., grid networks) regular networks (i.e., squared) compared to CRW. CRW remains around the neighborhood where it started expanding diffusively, whereas the superposition of QW produces a probability cloud expanding ballistically throughout the whole network (Kempe 2003; Venegas-Andraca 2012).

The superposition property of QW would theoretically allow a more efficient exploration process throughout the network, given the previously proposed conditions by McFadden and Al-Khalili (1999):

1. The cell is a quantum measurement device that constantly monitors the state of its own DNA molecule. The environment will induce the collapse of the quantum wave function, rendering the current state of the DNA (i.e., the DNA sequence we actually observe when we obtain the base pairs of a genome or a gene), indirectly via the influence of the environment on the cell (e.g., chemical conditions of the cell’s membrane and cytoplasm).
2. Following quantum mechanical jargon, the DNA molecule persists in a superposition of their hydrogen protons binding nucleotides (i.e., the different mutational options representing the wave function; see Godbeer et al. 2015). For instance, a wave function evolving to incorporate the correct and incorrect bases in a DNA position, as a superposition of states (i.e., mutated and unmutated states [e.g., the Cytosine and Thymine nucleotides in a DNA base pair]) in a daughter DNA strand; that is, the new DNA state achieved after replication of the genetic material (|**Ψ**_*G*_ >) (McFadden and Al-Khalili 1999):

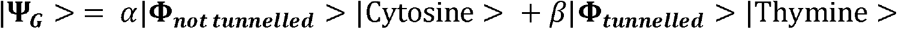
3. The operational difference between the DNA and the cell is given by nucleotides (previous equation above) and amino acids, respectively (see McFadden and Al-Khalili 1999):

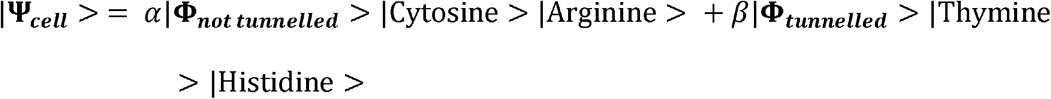
4. An evolving or new DNA wave function (i.e., the current DNA superposition after the collapse of the wave function due to environmental influences) must remain coherent or stable for long enough time to interact with the cell’s immediate environment, so the cell can act as a quantum device (Fig. 2).

### Genotype network construction

We used a subset of the DNA transcription factor genotype networks from the sample file of Genonets server (http://ieu-genonets.uzh.ch; Khalid et al. 2016), which represent empirical data for the binding affinities of the Ascl2, Foxa2, Bbx, and Mafb transcription factors (TF) in mice (Badis et al. 2009, Payne and Wagner 2014; Fig. 6). To filter genotypes with low binding affinities we used the default value of the parameter tau (τ = 0.35), and we only allowed for single point mutations (i.e., mutations where a letter in the sequence is changed, no indels were allowed; see http://ieu-genonets.uzh.ch/learn for definitions and tutorials; Khalid et al., 2016). Briefly, each node in the network represents a genotype with a specific TF phenotype (i.e., Ascl2, Foxa2, Bbx, Mafb), and the edges joining nodes represent mutational steps (i.e., two nodes joined by an edge are genotypes differing exactly by one position; in other words, only one mutation separates such nodes; see Figs. 3 and 4). We extracted the information of the genotype networks generated by Genonets, and performed all subsequent simulation analyses (described below) using the Mathematica software (Wolfram Research, Inc., 2020).

**Fig 6.**
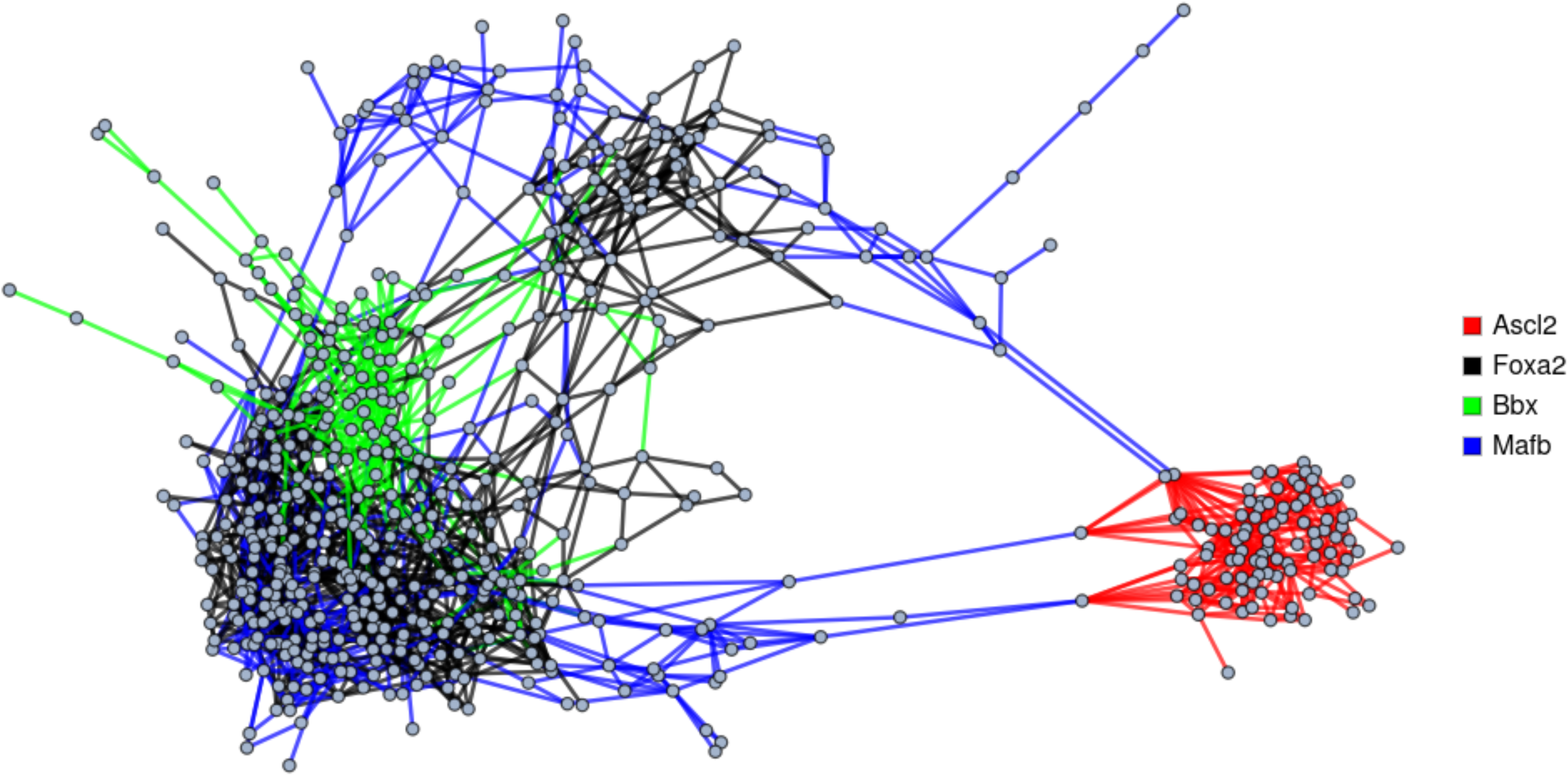
A subset of transcription factor genotype networks representing four phenotype networks (different colors) extracted from Genonets server (Khalid et al. 2016) and used for simulation analyses via QW and CRW.

### Genotype network exploration: closed systems (unitary evolution)

The n-dimensional genotype networks developed by Wagner and his collaborators use as an exploration mechanism CRW (Wagner 2011). Here, we used QW in order to explore the importance of quantum superposition (Farhi and Gutmann 1998; Mülken and Blumen 2011) as an evolutionary exploration device. Exploration of constructed networks was performed using both a continuous CRW (e.g., Rodrigues and Wagner 2009) and a continuous QW (Falloon et al., 2017). We used the QSWalk package developed under Mathematica to perform simulations on genotype networks (Falloon et al., 2017). The QSWalk package implements both CRW and continuous QW in arbitrary networks based on the so-called Quantum Stochastic Walk that generalizes quantum and classical random walks (Falloon et al., 2017).

For simplicity, we considered undirected and unweighted networks, which were described by an adjacency matrix **A**_ij_ whose matrix elements are 1 if the nodes i, j are connected and 0 otherwise. For an undirected network, the adjacency matrix is symmetric **A**_ij_ = **A**_ji_, which implies that transitions from any pair of neighboring nodes are equally probable independently of the direction. For each node i, we define the out-degree outDeg(j) = Σ_i ≠ j_ **A**_ij_, which counts the number of nodes connected to it. The CRW is described by the vector **p**(t) whose components p_j_(t) give the probability of occupancy of node j. The temporal evolution of the probability vector is determined by the equation

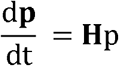

where **H** is the matrix

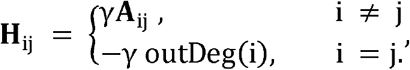

γ determines the transition rate between neighbor nodes. We considered CRW beginning in node i, implying that the components of the initial vector are p_k_(t = 0) = δ_{ki}_.

For the QW we considered a basis whose elements are associated to each node of the network | i >. A general pure state can be written as | Ψ(t)> = Σ_{i}_c_i_|i>, where |c_i_|^2^ is the probability of occupancy of node i. The dynamics of an initial configuration (similar to the CRW, we considered initial states with components given by c_k_ = δ_{ki}_) is given by the Schrödinger equation

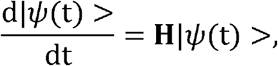

where **H** is a linear operator whose matrix elements are given by the same matrix introduced above

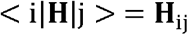

We used DNA transcription factor genotype networks with two (410 nodes) and four (927 nodes) different phenotypes, a representation of which is shown in Figure 6. Following, we determined the mutation rate, γ_c_ and γ_Q_, for the CRW and the QW respectively.

### Mutation rate, γ

The mutation rate between any pair of neighboring nodes is mapped in the QSWalk package by a parameter, γ. For a CRW, the probability of mutation of a given node to a new node for very short times is Pm = Nn × γ_c_ × t, where Nn is the number of neighboring nodes; in other words, the probability of remaining in the initial node decays exponentially. The average number of neighbor nodes in the networks used in our simulations is Nn ~ 6.4 ± 3.3; therefore the mutation rate per node is mr = 6.4 ± 3.3 γ_c_. Experimental estimation of this mutation rate (i.e., the rate of mutation of a single gene; Balin and Cascalho 2010) yields to mr = (4-9) × 10^−5^ mutations/base pair/cell generation. Assuming that a bacterial cell generation lasts around 1000 sec (i.e., ~20 minutes), the mutation rate is mr = (4-9) × 10^−8^ mutations/base pair/sec, which when equated to 6.4 ± 3.3 γ_c_ allows obtaining an estimation of the order of parameter γ_c_ for the CRW of γ_c_ = 10^−9^ ~ 10^−7^ (1/sec).

In contrast, for a quantum system described by a Hamiltonian, it can be shown that for very short times, the probability of transition of a given node to a new node grows quadratically with time (Mandelstam and Tamm 1945), Pm = Nn × (γ_Q_ × t)^2^. In order for the QW to be consistent with the experimental mutation rate mentioned above, we estimate γ_Q_, the mutation probability of a given node to a new node, by equating the quantum probability of node mutation with the classical probability at the decoherence time T_D_ (i.e., we considered γ_c_ × T_D_ = (γ_Q_ × T_D_)^2^, which gives γ_Q_^2^= γ_c_/ T_D_. Thus, to determine γ_Q_, an estimation of T_D_ is necessary. According to McFadden and Al-Khalili (1999), a rough estimation of the decoherence time is T_D_ =10^0^~10^2^ sec, which allows an estimation of quantum parameter γ_Q_ = 10^−6^ ~ 10^−3^. We selected representative values for γ_c_ and γ_Q_ to perform our simulations, γ_c_ = 10^−7^ (1/sec) and γ_Q_ = 10^−4^ (1/sec).

We follow McFadden and Al-Khalili (1999) at using the Zurek model to estimate the decoherence time of genotypes (nodes in the network) superposition (T_D_)

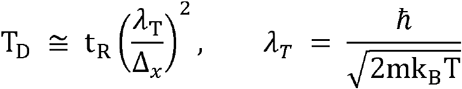

where m is the mass of a proton in a superposition of two Gaussian wave packets separated by a distance Δ_*x*_, and *λ*_*T*_ is the thermal de Broglie wavelength dependent (Djordjevic 2016).

For the small network consisting of 410 nodes and two phenotypes, we performed independent simulation runs with initial conditions that start from every single node of the Bbx phenotype to the Foxa2 phenotype and compared the probability to find the Foxa2 phenotype as a function of time for CRW and QW, distinguishing the number of mutational steps (1, 2 or 3) needed to reach the new phenotype.

For the larger network (927 nodes and 4 phenotypes), we conducted simulations starting at nodes that were shared between different pair-wise combinations of the four different phenotypic networks. The aim was to compare the efficiency at which CRW and QW find novel phenotypes as a function of time (for the quantum process within the decoherence time T_D_ as calculated above) and of the initial position of a node within a genotype network in terms of the number of mutational steps (i.e., 1, 2 or 3 edges) needed to reach the new phenotype.

## Results

### Two phenotype networks (Bbx and Foxa2; 410 nodes)

For this two-phenotype network, the linear dependence on time of the mutation probability of the CRW at short times, induces a linear dependence on time of the probability of mutation to phenotype Foxa2 from nodes located one mutational step away (Fig. 7). For nodes located two or three steps away, the growth of the mutation probability to the new phenotype is slower. The same hierarchy in the probabilities is observed in the QW, the closer the node is located to the new phenotype the larger the mutation probability to this phenotype is. Since the probability of an initial node to mutate to its neighboring nodes grows quadratically in the QW model, the probability of mutation to a new phenotype is smaller in the QW model for very short times. But at the temporal scale of quantum decoherence, the CRW and QW probabilities become comparable. Furthermore, for larger times, QW probabilities become much larger than the classical ones, irrespective of the distance of the initial node to the new phenotype. These results show that at a temporal scale comparable or slightly larger than the decoherence time, the QW becomes more efficient than the CRW at finding the new phenotype.

**Fig 7.**
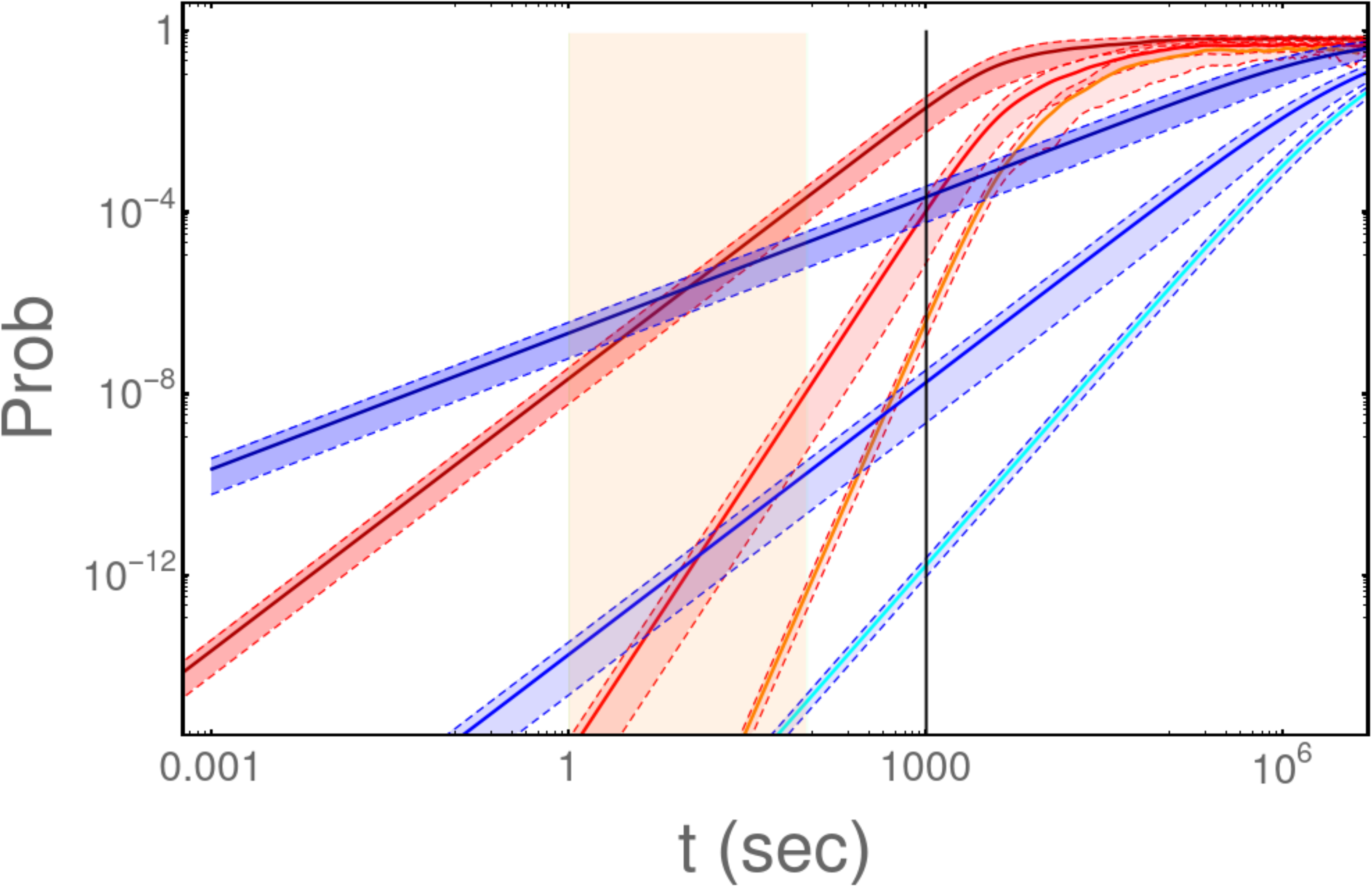
Simulation results for two phenotype networks (Foxa2 and Bbx; see Fig. 6). Probability of mutating to a novel phenotype as a function of time under the CRW (blue lines) and the QW (red lines) in log-log scale. Upper lines represent the average probability of simulations started at nodes that were one mutational step away from the novel phenotype; middle and lower lines are probability averages of nodes two and three mutational steps away from the novel phenotype, respectively. Shaded areas limited by dotted lines around the average lines represent the respective standard deviations of each simulation. The orange shaded area indicates the temporal estimates to decoherence time, and the vertical gray line is the time of a bacterial cell generation (i.e., approximately 20 minutes).

### Four phenotype networks (Ascl2, Foxa2, Mafb, and Bbx; 927 nodes)

Similar to the two-phenotype network simulation, for the CRW there is a linear dependency on the probability of mutating to a novel phenotype as a function of time (Fig. 8). For the QW at the temporal scale of quantum decoherence, the quadratic dependence of the probability of mutating makes the quantum process to have a higher probability of mutating to a novel phenotype under most conditions compared to the CRW. Such behavior was observed regardless of the number of mutational steps (i.e., 1, 2 or 3) away from the novel phenotype (Fig. 8), turning the QW effectively more efficient at finding novel phenotypes under different starting conditions. Furthermore, the QW became more efficient at finding novel phenotypes when the network increased in complexity in terms of the number of phenotypes and network size (compare QW results from Figs. 7 and 8).

**Fig 8.**
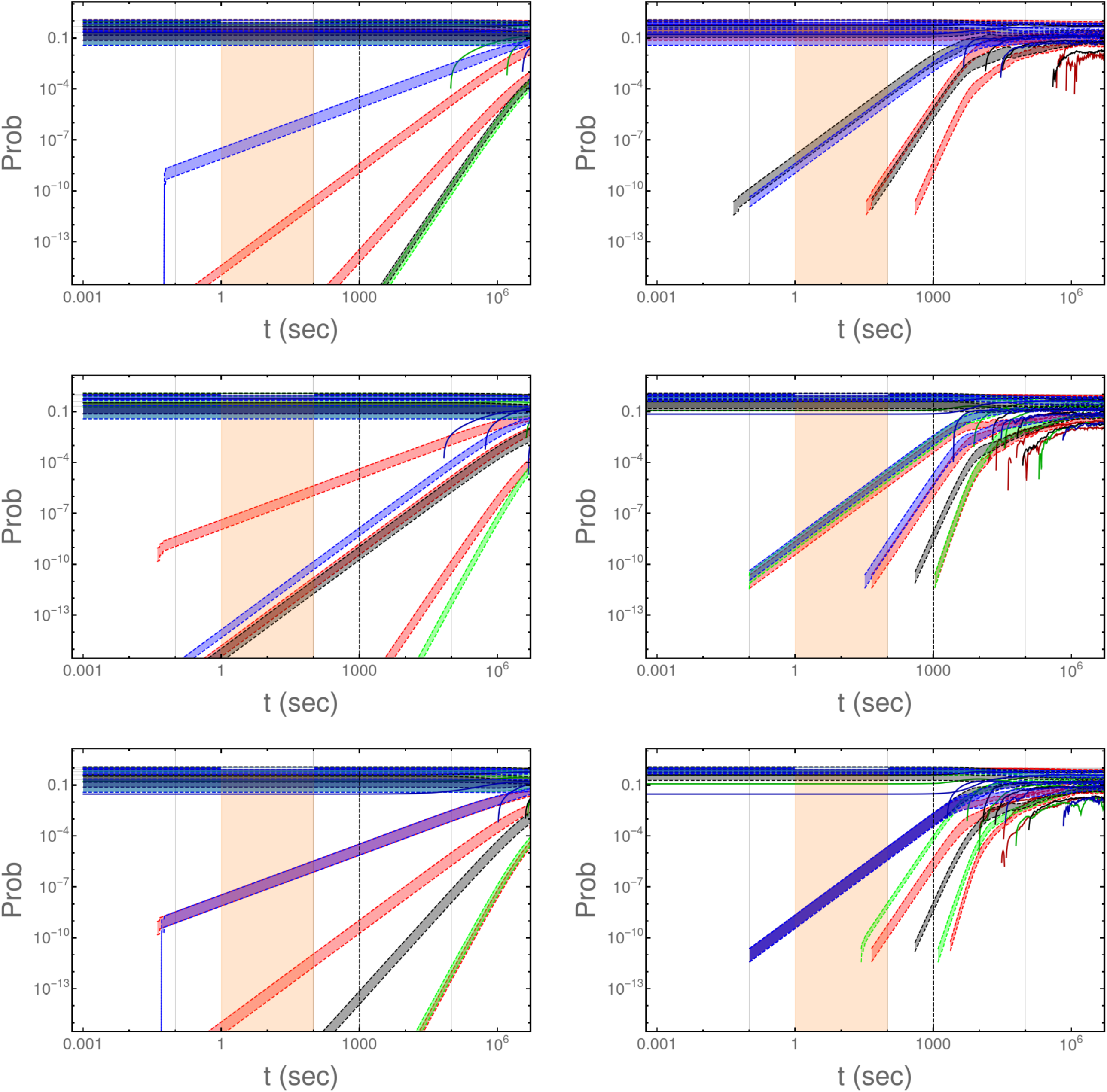
Simulation results for four phenotype networks (see Fig. 6). Probability of mutating to a novel phenotype as a function of time under the CRW (left column) and the QW (right column) in log-log scale. Top panels show results for nodes one mutational step away from a novel phenotype, middle panels for nodes two mutational steps away from novel phenotypes, and bottom panels started from three mutational steps away from novel phenotypes. The color of the different lines indicates the phenotype network where the simulation started (see Fig. 6). Shaded areas limited by dotted lines around the average lines represent the respective standard deviations of each simulation. Orange shaded areas indicate the temporal estimates to decoherence time, and the vertical dotted lines are the time of a bacterial cell generation (i.e., approximately 20 minutes).

## Discussion

The field of quantum biology has steadily grown over the last 15 years, in particular due to research focused on photosynthesis and enzymatic processes (Brooks 2017). However, advances on how quantum mechanisms are relevant to biological evolution have stagnated during the last two decades, most likely due to a lack of an evolutionary framework where such quantum processes can be studied (but see Martin-Delgado 2012; Asano et al. 2015). Here, we have suggested that n-dimensional genotype networks (*sensu* Wagner 2011) represent an ideal ground where the relevance of quantum superposition for evolution can be explored. We have shown that under neutral scenarios (i.e., non-selective environments or closed systems) QW become more efficient at the temporal scale of decoherence time and under more complex scenarios (four-phenotype vs. two-phenotype networks) than CRW. The QW model has exhibited a more diverse behavior in terms of mutation probabilities to a novel phenotype, which is readily observed under a varied array of conditions (i.e., when starting the simulations at 1, 2 or 3 mutational steps away from novel phenotypes). Interestingly, the efficiency of QW at finding novel phenotypes increased when the network structure increased in terms of number of phenotypes and size. This suggests that as network complexity (i.e., number of phenotypes) and size (number of genotypes or nodes) increases, we can expect the QW mechanism to be a more efficient exploration device for evolution given its superposition property. Thus, in order to move forward, the next step is to simulate QWs in open systems coupled to the environment, for example using dissipative Lindblad terms (e.g., Godbeer et al. 2015).

If QW prove indeed to be more efficient than CRW in an open network system, then the still controversial theoretical and experimental evidence in favor of adaptive mutations (e.g., Hall 1995, Palmer 2012, Braun and David 2011, Livnat 2013) would find an empirical framework supporting them. Of course, our proposal (i.e., quantum evolution on n-dimensional networks) does not preclude the existence and commonality of Darwinian random mutations; it only provides a complementary framework to understand currently suggested adaptive mutations. An example of a theory expanding current evolutionary understanding of mutations is that of the writing phenotype (Livnat 2013), which suggests that mutations are non-random in the sense that there is genomic data showing specific regions with higher rates of mutations due to specific genome structures. Mechanisms generating such non-random mutations include non-allelic homologous recombination, non-homologous DNA end-joining, replication-based mechanisms, and transposition (see Livant 2013 for details). In the cases of both n-dimensional genotype networks (Wagner 2009) and writing phenotypes (Livnat 2013) there are evolutionary constraints. In other words, non-random mutations (*sensu* Livnat 2013) are embedded in a genomic context that is modified as populations change from generation to generation. Hence, context dependent evolutionary constraints are dynamic because evolution shuffles the genomic context through time. However, such dynamic process does not necessarily mean that the procedure is blind to evolutionary direction within those constraints, which is where the quantum proposed mechanism of exploration on n-dimensional networks needs further study to determine its relevance. For example, by using dynamic adaptive networks.

### Philosophical extensions of QW to epigenetics and niche construction

Epigenetics investigates the regulatory mechanisms that during development lead to persistent and inducible heritable changes that do not affect the genetic composition of the DNA. Some of these changes can actually regulate the function of DNA without changing its base composition, via for example methyl groups (Jablonka and Lamb 2010). Epigenetic inheritance refers to those phenotypic variations that do not depend on DNA sequence variations, and that can be transmitted across generations of individuals (soma-to-soma) and cell lines (i.e., cellular epigenetic inheritance); such processes can lead to soft inheritance (Jablonka 2011). There are four basic types of epigenetic inheritance: 1) self-sustaining regulatory loops, where following the induction of gene activity, the gene’s own product acts as a positive feedback regulator maintaining gene’s activity across cell generations. 2) Structural templating, where preexisting 3D structures serve as models to build similar structures in the next generation of cells. 3) Chromatin markings, where small chemical groups (e.g., methyl CH_3_) bind to DNA, altering/controlling gene activity, they can segregate during DNA replication and be reconstructed in daughter DNA molecules. 4) RNA-mediated inheritance, where silent transcription states are maintained by interactions between small RNA molecules and their complementary mRNA and DNA. These states can be transmitted to cells and organisms via an RNA replication system, also by having small RNAs modifying heritable chromatin marks, and by inducing heritable gene deletions (Rassoulzadegan 2011; see Carey 2012 for a gently general introduction to epigenetics).

What is most relevant for the proposed framework is the fact that environmental factors (e.g., heat shock, starvation, chemicals, stress in general) can directly (germ line) or indirectly (somatic alterations) induce developmental modifications via heritable epigenetic variations, which underlie developmental plasticity and canalization (Nijhout 2003, Jablonka and Raz 2009, Jablonka 2011). If we implement the n-dimensional network concept of Wagner (2011) to an epi-genome, we can obtain an epigenetic network on which the environment can easily induce state changes in the expression and functioning of genes and even induce deletions and amplifications (Jablonka and Lamb 2010, see also Asano et al. 2015). Moreover, the response to the environment would be faster when less mutational or epi-mutational steps are required in reference to an environmental challenge (e.g., Blount et al. 2012). This last proposition can explain why in “clonal” bacterial evolutionary experiments not all colonies respond at the same time to the same environmental challenge, some respond differently but with similar results and some do not respond at all during the length of the experiment (e.g., Woods et al. 2006, Stanek et al. 2009, Braun and David 2011, Blount et al. 2012, Cooper 2012). The outcome will depend on exactly the structure of the genotype network and where on the genotype network evolution started during experiments.

Finally, niche construction is another non-Darwinian force imposing novel challenges on organisms via changes generated on the environment by the same organisms (Odling-Smee et al., 2003). In other words, changes imposed on the environment by species modify the adaptive landscape and the n-dimensional genotype network across generations. Such changes might produce environmental feedbacks on both the same organisms producing the change and indirectly on those other organisms under the influence of the novel environment. A novel environment will alter the probabilistic nature of the QW, changing the likelihood of evolutionary pathways (i.e., creating new evolutionary constraints), which according to our results would be better explored by the diverse behavior of the QW than CRW.

The framework presented here provides a probabilistic process (via a quantum wave function) that might act as the mechanism for the evolutionary exploration of n-dimensional genotype networks within the constraints established by the available options (i.e., phenotypes). In this sense, our study complements the initial work of Ogryzko (1997) and McFadden and Al-Khalili (1999) by providing an evolutionary context (highly diverse and robust n-dimensional genotype networks), where a quantum wave function is the mechanism of evolutionary exploration. The process still needs to be investigated in much larger n-dimensional genotype networks and also under open system scenarios, where the environment might influence system’s behavior. Such analysis will determine if certain cell states (quantum superposition) have stronger interactions with current environmental conditions compared to other states, which subsequently promotes quantum decoherence toward those more likely options resulting in adaptive mutations (e.g., Asano et al. 2015; Godbeer et al. 2015). Those likely options will be given by the current genomic context of the population (i.e., the n-dimensional genotype network), which are not necessarily better or best for the current conditions, but are most likely in accordance to current context (i.e., evolutionary constraints; see Rozen et al. 2008 for a probable example of this effect).

A way to prove our theory experimentally can be by using clonal bacterial colonies that start from different positions in the genotype network, in such a way that decoherence times can be measured under the influence of a novel environment (e.g., lactose); such times should be repeatable across experiments (see Fig. 5). Modern –omics (e.g., genomics, transcriptomics) and biotechnology techniques can be used to construct specific bacterial lines for such experiments. In addition, it would be possible to analyze the epi-genome of plants, which are the organisms where this type of non-Darwinian evolutionary process is more common.

## Supporting information

Building genotype networks code

Calculating probabilities for CRW and QW

Simulations for Classical Random Walks

Simulations for Quantum Walks

Mathematica notebook build network

Mathematica notebook probabilities

Mathematica notebook simulation CRW

Mathematica notebook simulation QW

## Acknowledgements

We are grateful to A. Raúl Hernández Montoya for providing computational resources. DS-A has been continuously supported by the Consejo Nacional de Ciencia y Tecnología de México (CONACyT project number Ciencia Básica 2011-01-168524 and project number Problemas Nacionales 2015-01-1628) and the Instituto de Ecología, A.C. S.L.-H. acknowledges financial support from CONACYT project CB2015-01/255702. SEV-A acknowledges the financial support of Tecnológico de Monterrey, Escuela de Ingeniería y Ciencias and of CONACyT (SNI number 41594, as well as Fronteras de la Ciencia project No. 1007).

## Data, Code and Materials

The Mathematica code and data used for simulations supporting results of this article are available upon request to and will be publicly stored in figshare.com once the paper is published. See also supplementary material.

## Authors’ contributions

DS-A developed the idea and drafted the manuscript; DS-A, HT-Mc, and SL-H refined the concept and designed the simulations; HT-Mc and SL-H performed the simulations and helped draft the manuscript. SEV-A refined the idea, formalized the quantum random walks mathematics, and helped draft the manuscript. All authors gave final approval for publication and agreed to be held accountable for the work performed therein.

## References

Aguilar-Rodríguez J, Payne JL, Wagner A. 2017. A thousand empirical adaptive landscapes and their navigability. Nature Ecology & Evolution 1: 0045.

Asano M, Khrennikov A, Ohya M, Tanaka Y, Yamato I. 2015. Quantum adaptivity in biology: from genetics to cognition. Springer, Dordrecht

Badis G, Berger MF, Philippakis AA, Talukder S, Gehrke AR, Jaeger SA, Chan ET, Metzker G, Vedenko A, Chen X, Kuznetsov H, Wang CF, Coburn D, Newburger DE, Morris Q, Hughes TR, Bulyk ML. 2009. Diversity and complexity in DNA recognition by transcription factors. Science 324: 1720–1723.

Balin SJ, Cascalho M. 2010. The rate of mutation of a single gene. Nucleic Acids Research. 38: 1575–1582.

Bastolla U, Porto M, Roman HE, Vendruscolo M. 2003. Connectivity of neutral networks, overdispersion, and structural conservation in protein evolution. Journal of Molecular Evolution 56: 243–254.

Blount ZD, Barrick JE, Davidson CJ, Lenski RE. 2012. Genomic analysis of a key innovation in an experimental *Escherichia coli* population. Nature 489: 513–518.

Braun E, David L. 2011. The role of cellular plasticity in the evolution of regulatory novelty. In Transformations of Lamarckism: from subtle fluids to molecular biology (Gissis, S.B. and E. Jablonka, Editors). MIT Press. Massachusetts, USA. Pp. 181–191.

Brovarets’ OO, Hovorun DM (2015) Proton tunneling in the A·T Watson-Crick DNA base pair: myth or reality?, Journal of Biomolecular Structure and Dynamics, 33: 2716–2720.

Brookes JC. 2017. Quantum effects in biology: golden rule in enzymes, olfaction, photosynthesis and magnetodetection. Proceedings of the Royal Society A 473: 20160822.

Cairns J, Overbaugh J, Millar S. 1988. The origin of mutants. Nature 335: 142–145.

Carey, N. 2012. The epigenetics revolution. Columbia University Press, New York.

Ciliberti S, Martin OC, Wagner A. 2007. Robustness can evolve gradually in complex regulatory gene networks with varying topology. PLoS Computational Biology 3: e15.

Cooper TF. 2012. Empirical insights into adaptive landscapes from bacterial experimental evolution. In The adaptive landscape in evolutionary biology (Svensson, E.I., and Calsbeek, R., Editors). Oxford Univ. Press. Oxford, U.K. Pp. 169–179.

Cui Y, Wong W, Bornberg-Bauer E, Chan H. 2002. Recombinatoric exploration of novel folded structures: a heteropolymer-based model of protein evolutionary landscapes. Proceedings of the National Academy of Sciences USA 99: 809–814.

Djordjevic IV. 2016. Quantum biological information theory. Springer, Cham.

Espinosa-Soto C, Martin OC, Wagner A. 2011. Phenotypic robustness can increase phenotypic variability after nongenetic perturbations in gene regulatory circuits. Journal of Evolutionary Biology 24: 1284–1297.

Falloon PE, Rodriguez J, Wang JB. 2017. QSWalk: a Mathematica package for quantum stochastic walks on arbitrary graphs. Computer Physics Communications 217: 162–170.

Farhi E, Gutmann S. (1998). Quantum computation and decision trees. Physical Review A 58: 915–929.

Godbeer AD, Al-Khalili JS, Stevenson PD. 2015. Modelling proton tunneling in the adenine-thymine base pair. Physical Chemistry Chemical Physics 17: 13034.

Hall BG. 1995. Adaptive mutations in *Escherichia coli* as a model for the multiple mutational origins of tumors. Proceedings of the National Academy of Sciences USA 92: 5669–5673.

Hall BG. 1997. Spontaneous point mutations that occur more often when advantageous than when neutral. Genetics 126: 5–16.

Hall BG. 1998. Adaptive mutagenesis at ebgR is regulated by PhoPQ. Journal of Bacteriology 180: 2862–2865.

Hendrickson H, Slechta ES, Bergthorsson U, Andersson DI, Roth JR. 2002. Amplification-mutagenesis: evidence that “directed” adaptive mutation and general hypermutability result from growth with a selected gene amplification. Proceedings of the National Academy of Sciences USA 99: 2164–2169.

Jablonka E, Raz G. 2009. Transgenerational epigenetic inheritance: prevalence, mechanisms, and implications for the study of heredity and evolution. Quarterly Review of Biology 84: 131–176.

Jablonka E, Lamb MJ. 2010. Transgenerational epigenetic inheritance In Evolution the extended synthesis (Pigliucci, M. and Müller, G.B., Editors). MIT Press. Cambridge, MA. USA. Pp. 137–174.

Jablonka E. 2011. Cellular epigenetic inheritance in the twenty-first century. In Transformations of Lamarckism: from subtle fluids to molecular biology (Gissis, S.B. and E. Jablonka, Editors). MIT Press. Massachusetts, USA. Pp. 215–226.

Kempe J (2003) Quantum random walks: An introductory overview, Contemporary Physics 44: 307–327.

Khalid F, Aguilar-Rodríguez J, Wagner A, Payne JL 2016. Genonets server – a web server for the construction, analysis and visualization of genotype networks. Nucleic Acids Research 44: W70–W76.

Klinman JP, Kohen A. 2013 Hydrogen tunneling links protein dynamics to enzyme catalysis. Annual Review of Biochemistry 82: 471–496.

Li H, Helling R, Tang C, Wingreen N. 1996. Emergence of preferred structures in a simple model of protein folding. Science 273: 666–669.

Livnat A. 2013. Interaction-based evolution: how natural selection and nonrandom mutation work together. Biology Direct 8: 24.

Lloyd S. 2008. Quantum mechanics and emergence. In Quantum aspects of life (Abbott, D., Davies, P.C.W., and Pati, A.K., Editors). Imperial College Press. London, UK., pp. 19–30.

Mandelstam L, Tamm I. 1945. The uncertainty relation between energy and time in nonrelativistic quantum mechanics. Journal of Physics (USSR) 9:249–254.

Martin-Delgado JA. 2012. On quantum effects in a theory of biological evolution. Sci. Rep. 2:302. DOI: 10.1038/srep00302

McFadden J, Al-Khalili J. 1999. A quantum mechanical model of adaptive mutation. BioSystems 50: 203–211.

McFadden J, Al-Khalili J. 2014. Life on the edge: the coming of age of quantum biology. Crown Publishers. New York, NY.

McFadden J, Al-Khalili J. 2018 The origins of quantum biology. Proceedings of the Royal Society of London A 474: 20180674.

Mohseni M, Omar Y, Engel G, Plenio M (Eds.). 2014. Quantum effects in biology. Cambridge University Press. Cambridge, UK.

Mülken O, Blumen A. 2011. Continuous-time quantum walks: models for coherent transport on complex networks. Physics Reports 502:37–87.

Nijhout HF. 2003. Development and evolution of adaptive polyphenisms. Evolutionarly Development. 5: 9–18.

Odling-Smee FJ, Laland KN, Feldman MW. 2003. Niche Construction: the neglected process in evolution. Princeton Univ. Press. Princeton, NJ, USA.

Ogryzko VV. 1997. A quantum-theoretical approach to the phenomenon of directed mutations in bacteria (hypothesis). BioSystems 43: 83–95.

Palmer AR. 2012. Developmental plasticity and the origin of novel forms: unveiling cryptic genetic variation via “use and disuse”. Journal of Experimental Zoology B: Molecular and Developmental Evolution 318: 466–479.

Panitchayangkoon G, Hayes D, Fransted KA, Caram JR, Harel E, Wen J, Blankenship RE, Engel GS. 2010. Long-lived quantum coherence in photosynthetic complexes at physiological temperature. Proceedings of the National Academy of Sciences USA 107: 12766–12770.

Payne JL, Wagner A. 2014. The robustness and evolvability of transcription factor biding sites. Science 343: 875–877.

Rassoulzadegan M. 2011. An evolutionary role for RNA-mediated epigenetic variation? In Transformations of Lamarckism: from subtle fluids to molecular biology (Gissis, S.B. and E. Jablonka, Editors). MIT Press. Massachusetts, USA. Pp. 227–235.

Rodrigues JFM, Wagner A. 2009. Evolutionary plasticity and innovations in complex metabolic reaction networks. Plos Computational Biology 5:e1000613.

Rodrigues JFM, Wagner A. 2011. Genotype networks, innovation, and robustness in sulfur metabolism. BMC Systems Biology 5:39

Rosenberg SM, Longerich S, Gee P, Harris RS. 1994. Adaptive mutation by deletions in small mononucleotide repeats. Science 265: 405–407.

Rozen DE, Habets MGJL, Handel A, de Visser JAGM (2008) Heterogeneous Adaptive Trajectories of Small Populations on Complex Fitness Landscapes. PLoS ONE 3(3): e1715.

Samal A, Rodrigues JFM, Jost J, Martin OC, Wagner A. 2010. Genotype networks in metabolic reaction spaces. BMC Systems Biology 4:30.

Samal A, Wagner A, Martin OC. 2011. Environmental versatility promotes modularity in genome-scale metabolic networks. BMC Systems Biology 5:135.

Schrödinger E. 1944. What is Life? Cambridge University Press, London.

Schuster P, Fontana W, Stadler P, Hofacker I. 1994. From sequences to shapes and back – a case-study in RNA secondary structures. Proceedings of the Royal Society of London B 255: 279–284.

Stanek MT, Cooper TF, Lenski RE. 2009. Identification and dynamics of a beneficial mutation in a long-term evolution experiment with *Escherichia coli*. BMC Evolutionary Biology 9: 302.

Stumpf JD, Poteete AR, Foster PL. 2007. Amplification of *lac* cannot account for adaptive mutation to Lac^+^ in *Escherichia coli*. Journal of Bacteriology 189:2291–2299.

Sumedha, Martin, OC, Wagner A. 2007. New structural variation in evolutionary searchers of RNA neutral networks. Biosystems 90: 475–485.

Venegas-Andraca SE. 2012. Quantum walks: a comprehensive review. Quantum Information Processing 11:1015–1106.

Wagner A. 2008. Robustness and evolvability: a paradox resolved. Proceedings of the Royal Society of London B 275: 91–100.

Wagner A. 2009. Evolutionary constraints permeate large metabolic networks. BMC Evolutionary Biology 9: 231.

Wagner A. 2011. The origins of evolutionary innovations: a theory of transformative change in living systems. Oxford Univ. Press. New York. USA.

Wagner A. 2012. The role of randomness in Darwinian evolution. Philosophy of Science 79:95–119.

Wagner A. 2014. Arrival of the fittest: solving evolution’s greatest puzzle. Current. New York, USA.

Watson JD, Crick FHC. 1953a. Molecular structure of nucleic acids: a structure for deoxyribose nucleica cid. Nature 171: 737–738.

Watson JD, Crick FHC. 1953b. Genetical implications of the structure of deoxyribonucleic acid. Nature 171: 964–967.

Wolfram Research, Inc. 2020. Mathematica, v12.1, Champaign, USA.

Woods R, Schneider D, Winkworth CL, Riley MA, Lenski RE. 2006. Tests of parallel molecular evolution in a long-term experiment with *Escherichia coli*. Proceedings of the National Academy of Sciences USA 103: 9107–9112.

