## Supplementary material for "Quantum aspects of evolution: a contribution toward evolutionary explorations of genotype networks via quantum walks": Building genotype networks code

### Supplementary notebook 1: Build the genotype network.

- This notebook processes the source file downloaded from the Genonets Server: http://ieu-genonets.uzh.ch/
- Source file is **“`genonetsT1.json.gz`”** provided also as supplementary material. It must be placed in a subdirectory inside the path where the notebook is evaluated. Otherwise change to the correct path in the variable **`$dataURL`**
- Results of this notebook’s evaluation are two structures with the network’s information (also provided as supplementary data):
  - **“`g.wl`”** is an Association containing the nodes and its connections (genes and targets)
  - **“`gN.wl`”** is a list of undirected edges used to represent the graph object
- The two structures are used to generate the classical and quantum walks in another notebook also provided as supplementary material.
- **It is not necessary to evaluate this notebook as the results of its evaluation are also provided as supplementary electronic material.**

```
ClearAll["Global`*"]
SetOptions[$FrontEndSession, NotebookAutoSave → True]
NotebookSave[]
$baseURL = DirectoryName[NotebookDirectory[]];
$dataURL = FileNameJoin[{$baseURL, "Data"}];
```

In[ ]:=
```
Clear[makeGraphFromSequence]
makeGraphFromSequence[genet_] :=
 Module[{vertices, edgesAsIndexPairs, edges, evol2, evolTargets, evolEdges},
  vertices = genet[["nodes", All, "sequences"]];
edgesAsIndexPairs = 1 + Values /@ genet[["edges", All, {"source", "target"}]];
edges = UndirectedEdge @@ vertices[[#]] & /@ edgesAsIndexPairs;
  evol2 = genet[["nodes", All, "Evolvability_targets"]];
  evolTargets = Values /@ evol2;
  evolEdges = UndirectedEdge @@@ Flatten[Table[
      Outer[List, {vertices[[n]]}, evolTargets[[n]]], {n, Length[vertices]}
     ], 3];
  (*{vertices,Join[edges,evolEdges]}*)
  <|"Genes" → vertices, "Targets" → Join[edges, evolEdges]|>
 ]
```

In[ ]:=
```
netdata = Import[FileNameJoin[{$dataURL, "genonetsT1.json.gz"}], "RawJSON"];
Position[Values@netdata[[ ;; , "graph", "Evolvability"]], _Missing]
Extract[Keys@netdata, %]
netdata = Delete[netdata, First@%];
```

In[ ]:=
```
g = makeGraphFromSequence /@ netdata;
phenotypes = Keys@g;
```

In[ ]:=
```
gN = g[#, "Targets"] & /@ phenotypes;
```

In[ ]:=
```
geneTargets = Values[EdgeRules /@ gN];
partGeneTargets = Part[geneTargets, #] & /@ Subsets[Range[pheNLen], {2}];
commonGenes = DeleteCases[Intersection @@ # & /@ partGeneTargets, {}];
commGenes = Intersection @@ # & /@ partGeneTargets;
Length /@ {commonGenes, commGenes}
```

```
Save[FileNameJoin[{$dataURL, "gN.wl"}], gN]
Save[FileNameJoin[{$dataURL, "g.wl"}], g]
```
