## Supplementary material for "Quantum aspects of evolution: a contribution toward evolutionary explorations of genotype networks via quantum walks": Calculating probabilities for CRW and QW

### Supplementary Notebook 2: Simulation of the genotype network.

- This notebook calculates the classical and quantum probabilities of evolving from a genotype on a phenotype to another using the functionality provided in the package `QSWalk` available in https://doi.org/10.1016/j.cpc.2017.03.014
- The full network is stored in variable **“G”** created in this notebook, from which the generator matrix is built.
- ***n*** number of vertex count on graph
- **neig** gives the list of vertex on the neighborhood graph common to genotypes for a pair of phenotypes
- **neigborsIdx** gives the index on the full genotype network of these neighbor genotypes
- **particiones** contains the index of the genotypes on the full network
- **dtes** contains the times for the simulation
- **It is not necessary to evaluate this notebook as the results are also provided as supplementary electronic material.**
- Note that the calculations performed by this code are memory intensive and the resulting files are large (2 - 4 Gb)
- The results of these evaluations are stored in the provided data files named **`classicalProbsUnPasoQuePartenDe####.wl`** and **`quantumProbsUnPasoQuePartenDe####.wl`**, where #### is the name of the phenotype network from where the calculations begin.

### Initialization: import data, build network and functions to perform the classical and quantum walks

### For each target network, identified by the index p the following will calculate classical and quantum probabilities

```
p = 1; (*"Ascl2"→"Foxa2"*)
genoTargets[[p]]
neighbors = VertexList[NeighborhoodGraph[G, commonGenes[[p]]]]
neighborsIdx = VertexIndex[G, #] & /@ neighbors
```

```
Monitor[
 classicalProbfromNeighbors2All =
   Table[calculaProbabilidadClasica[pi, dtes, stc, #], {pi, neighborsIdx}] & /@
    particiones;,
 pi]
```

```
Monitor[
 quantumProbfromNeighbors2All =
   Table[calculaProbabilidadCuantica[pi, dtes, stq, #], {pi, neighborsIdx}] & /@
    particiones;,
 pi]
```
