## Supplementary material for "Quantum aspects of evolution: a contribution toward evolutionary explorations of genotype networks via quantum walks": Simulations for Classical Random Walks

#### Supplementary Notebook 3: Data analysis of classical random walk simulations

- This notebook plots the **classical** probabilities computed using the functionality provided by the **QSWalk** package available in https://doi.org/10.1016/j.cpc.2017.03.014
- The timings shown on this notebook where obtained on a LENOVO ThinkStation E31 tower computer equipped with 4 64 bit Intel(R) Core(TM) i5-3450 CPU @ 3.10GHz and 32GiB of RAM running Ubuntu 18.04.4 LTS
- The inputs are provided as supplementary material with names under a directory tree with the following structure: (this notebook reads the data in the directory **classical**)

##### Directory tree structure

### Classical Un steps from mutation

In[ ]:=
```
qPath = FileNames[All, FileNameJoin[{path, "UnPaso", "classical"}]]
```

#### First dataset, Ascl2upq

In[ ]:=
```
AbsoluteTiming[
 Ascl2upq = Get[qPath[[1]]];
]
Print["Took ", %[[1]] / 60., " minutes to read data"]
```

Out[ ]= {240.08, Null}

```
Took 4.00133 minutes to read data
```

Iterate manually the code below to create the plt# for each phenotype network

```
In[ ]:= Table[
  dts = Ascl2upq[[n, ;; , 1]][[1, All, 1]];
  ave = Mean[Ascl2upq[[n, ;; , 1]][[All, All, 2]]];
  std = StandardDeviation[Ascl2upq[[n, ;; , 1]][[All, All, 2]]];

  Ascl2upqPlot[n] = ListPlot[Transpose[{dts, #}] & /@ {ave - std, ave, ave + std},
    Filling -> {2 → {{3}, {Opacity[0.35, Red]}}},
    PlotStyle → {{Thick, Darker[Red]}, {Red, Dashed}, {Red, Dashed}},
    PlotRange → All, GridLines → Automatic, Joined → True(*,
    PlotLabel→keys[[p]]*), ScalingFunctions → {"Log", "Log"}];, {n, 4}
 ]
Out[ ]= {Null, Null, Null, Null}

In[ ]:= qPlotAscl2 = Show[Ascl2upqPlot /@ Range[4], re, rec, Frame → True, Axes → False,
    FrameStyle → Thick, FrameTicksStyle → Large, PlotRange → {{-7, 15}, {-35, 0.7}},
    FrameLabel → {Style["t (sec)", 40], Style["Prob", 40]}, ImageSize → 800];
 Export[fileName["Un", "Ascl2"], %]
Out[ ]= ClassicalProbsUnPasoDeAscl2 .png

In[ ]:= fileName["Un", "Ascl2"]
Out[ ]= ClassicalProbsUnPasoDeAscl2 .tiff
```

##### Second dataset, Bbxupq

##### Third dataset, Foxa2upq

##### Fourth dataset, Mafbupq

##### Combine plots

### Classical Dos steps from mutation

### Classical Tres steps from mutation
