## Supplementary material for "Quantum aspects of evolution: a contribution toward evolutionary explorations of genotype networks via quantum walks": Simulations for Quantum Walks

#### Supplementary Notebook 3: Data analysis of quantum walk simulations

- This notebook plots the **quantum** probabilities computed using the functionality provided by the **QSWalk** package available in https://doi.org/10.1016/j.cpc.2017.03.014
- The timings shown on this notebook where obtained on a LENOVO ThinkStation E31 tower computer equipped with 4 64 bit Intel(R) Core(TM) i5-3450 CPU @ 3.10GHz and 32GiB of RAM running Ubuntu 18.04.4 LTS
- The files are provided in a directory tree with the following structure: (this notebook reads the data in the directory **quantum**)

##### Directory tree structure

### Quantum Un steps from mutation

```
In[ ]:= qPath = FileNames[All, FileNameJoin[{path, "UnPaso", "quantum"}]];
```

#### First dataset, Ascl2upq

```
In[ ]:= AbsoluteTiming[
         Ascl2upq = Get[qPath[[1]]];
        ]
        Print["Took ", %[[1]] / 60., " minutes to read data"]

Out[ ]= {785.351, Null}

        Took 13.0892 minutes to read data
```

Iterate manually the code below to create the plt# for each phenotype network

In[ ]:=
```
Table[
 dts = Ascl2upq[[n, ;; , 1]][[1, All, 1]];
 ave = Mean[Ascl2upq[[n, ;; , 1]][[All, All, 2]]];
 std = StandardDeviation [Ascl2upq[[n, ;; , 1]][[All, All, 2]]];

 Ascl2upqPlot [n] = ListPlot[Transpose [{dts, #}] & /@ {ave - std, ave, ave + std},
   Filling -> {2 → {{3}, {Opacity[0.35, Red]}}},
   PlotStyle → {{Thick, Darker[Red]}, {Red, Dashed}, {Red, Dashed}},
   PlotRange → All, GridLines → Automatic, Joined → True(*,
   PlotLabel→keys[[p]]*), ScalingFunctions → {"Log", "Log"}];, {n, 4}
]
```

Out[ ]=
```
{Null, Null, Null, Null}
```

In[ ]:=
```
qPlotAscl2 = Show[Ascl2upqPlot /@ Range[4], re, rec, Frame → True, Axes → False,
   FrameStyle → Thick, FrameTicksStyle → Large, PlotRange → {{-7, 15}, {-35, 0.7}},
   FrameLabel → {Style["t (sec)", 40], Style["Prob", 40]}, ImageSize → 800];
Export[fileName["Un", "Ascl2"], %]
```

Out[ ]=
```
QuantumProbsUnPasoDeAscl2 .png
```

##### Second dataset, Bbxupq

##### Third dataset, Foxa2upq

##### Fourth dataset, Mafbupq

##### Combine plots

#### Quantum Dos steps from mutation

#### Quantum Tres steps from mutation
